# The SWI/SNF complex sub-unit Bap60 is required for training-induced gene transcription during long-term memory formation

**DOI:** 10.1101/2025.08.24.671976

**Authors:** Spencer G. Jones, Nicholas Raun, Shanyn C. Bleeker, Abigail L. Henn, Jamie M. Kramer

**Affiliations:** Biochemistry and Molecular Biology, Dalhousie University, Halifax, Canada

**Keywords:** Long-term memory formation, *Drosophila melanogaster*, transcription, chromatin, Bap60, SMARCD1, SWI/SNF

## Abstract

Long-term memory (LTM) formation requires tightly regulated gene expression in response to neuronal activity. Chromatin remodeling is thought to play a role in enabling this transcriptional response, but the specific mechanisms remain unclear. In *Drosophila*, a transcriptional trace of courtship LTM training can be observed in the mushroom body (MB) during the memory consolidation phase after the end of training. Here, we investigated the role of Bap60, a core subunit of the *Drosophila* SWI/SNF chromatin remodeling complex, in the transcriptional trace of memory consolidation. Adult-specific knockdown of *Bap60* in the MB selectively impaired LTM with short-term memory (STM) left intact. Transcriptome analysis of Bap60 knockdown MBs following courtship LTM training revealed near complete disruption of LTM training induced transcription in the MB. MB-specific CUT&RUN was used to map SWI/SNF binding sites and identify over 100 candidate direct Bap60-dependent training induced genes. Bap60 was not required for training induced activation of immediate early genes (IEGs), Hr38 and Sr, which are transcription factors that are critical for courtship LTM. This suggests that Bap60 may regulate LTM gene induction downstream of IEGs. Interestingly, we identified Sr binding sites at 30% of Bap60-dependent training induced genes. Many of these target genes were transcription factors, including Prospero (Pros), which we show is also required in the MB for LTM but not STM. Interestingly, Pros is induced in a subset of MB nuclei, suggesting Bap60 is involved in cell specific LTM training induced gene activation. Together, these findings reveal a critical role for Bap60, a core SWI/SNF subunit, as a key regulator of LTM training-induced transcription downstream of IEGs.

**Significance Statement:** Autosomal dominant mutations in SWI/SNF subunits are a leading cause of neurodevelopmental disorders (NDDs) including Autism and Intellectual Disability. Therefore, understanding their role in brain function can provide insight into the mechanisms underlying human disease. The SWI/SNF complex is an important chromatin remodeling complex that is conserved from yeast to mammals. This large protein complex is a transcriptional coactivator that uses energy from ATP to open up chromatin and create binding sites for transcription factors. Our knowledge about how SWI/SNF functions in different cell types of the brain is highly limited. In this study we investigate the Bap60 component of SWI/SNF in memory neurons of the fruit fly, *Drosophila melanogaster*. We find that Bap60 is required for activation of gene expression programs underlying long term memory consolidation. Dynamic gene expression, like that associated with LTM, is essential to build neural circuits during human cognitive development. It is therefore possible that this process is disrupted in people with SWI/SNF-related NDDs.

## Introduction

Long-term memory (LTM) formation requires a transcriptional response to experience that produces new proteins critical for stabilizing memory traces^1^. This response involves the induction of immediate early genes (IEGs), whose expression is induced by neuronal activity^2,3^. Across diverse species and memory paradigms, transcription factors such as CREB initiate early waves of IEG expression at the onset of memory formation^4–12^. However, the downstream effectors and the chromatin-associated mechanisms that regulate the memory-relevant transcriptional response remain incompletely defined.

The SWI/SNF complex is an evolutionarily conserved ATP-dependent chromatin remodeler first identified in yeast and present throughout metazoans^13–15^. In mammals, this complex is comprised of approximately 15 subunits encoded by 29 genes, many of which harbor variants linked to neurodevelopmental disorders, including Autism Spectrum Disorders and Intellectual Disability^15,16^. SWI/SNF modulates DNA accessibility by repositioning nucleosomes and thereby enables transcription factors to engage promoters and enhancers and regulate gene expression^17–20^. While the roles SWI/SNF plays in cell fate specification and neuronal differentiation during development are well established^21–26^, SWI/SNF-mediated gene regulation in post-mitotic neurons is less well explored. Interestingly, in mammals there is a neuron specific SWI/SNF complex called nBAF, that contains the Baf53B subunit only in postmitotic neurons^27–30^. Baf53B is required for neuron activity induced dendritic growth in cell culture, and synaptic plasticity and memory in adult mice^20,27^. However, there are conflicting results about the role of Baf53B in regulating IEG expression^28^. The potential role of other SWI/SNF subunits in neuron activity induced gene expression has not been explored.

We recently described a transcriptional trace of LTM using *Drosophila* courtship conditioning^31^, a memory assay where male flies are trained by prolonged exposure to a non-receptive mated female that rejects copulation attempts^32,33^. As a result, trained males learn to suppress courtship behavior toward unreceptive females, and this suppression can be quantitatively measured during subsequent encounters^34,35^. Courtship LTM is dependent on the mushroom body (MB)^36^, which is the general memory center of the fly brain^37–40^. We have shown that memory genes are upregulated in the MB 1 hour after LTM training, during the memory consolidation phase^31,41,42^. Here, we investigate the *Drosophila* SWI/SNF subunit Bap60 and its role in the LTM training induced transcriptome.

Bap60 is the *Drosophila* ortholog of SMARCD1, a core SWI/SNF subunit. Autosomal dominant *SMARCD1* mutations cause a syndromic neurodevelopmental disorder characterized by Intellectual Diability^43^. We show that post-developmental knockdown of *Bap60* selectively impairs LTM, without effecting STM. MB cell-type specific transcriptome profiling following courtship training revealed defective induction of training-induced gene transcription, including genes associated with memory. We mapped SWI/SNF MB targets and identified the transcription factor Prospero (Pros) as a candidate memory regulator downstream of Bap60. Overall, these results establish Bap60 as a critical regulator of LTM training induced transcription in the MB.

## Results

### Adult-specific knockdown of Bap60 in the MB abolishes LTM, but not STM

To investigate the role of the SWI/SNF chromatin remodeling complex in acute memory processes, we performed adult-specific RNAi knockdown of the core subunit Bap60 in MB neurons using the *R14H06-Gal4* driver, which is specifically expressed in post-mitotic MB neurons, including MBα/β and MBγ neurons^31,41,44^. This driver was used to express a previously validated^43,45^ RNAi construct targeting *Bap60*, alongside *5xUAS-unc84::2xGFP*, a *Caenorhabditis elegans*-derived nuclear tag enabling used for Isolation of Nuclei in a Tagged Cell Type (INTACT)^46^. Expression was controlled by temperature-sensitive *Gal80* (*Gal80^ts^*) to restrict knockdown to adult flies. Previously we induced Bap60 knockdown (Bap60-KD) in newly eclosed adults and this caused defects in both STM and LTM^43^. Here, we initiated knockdown 4 days after eclosion, one day prior to training **(Fig. 1A)**. Under these conditions, Bap60-KD flies were able to form STM, displaying significantly reduced courtship activity one hour after training compared to naïve flies, and overall STM was significantly enhanced relative to genetic controls **(Fig. 1A)**. In contrast, Bap60-KD flies failed to retain courtship suppression 24 hours after LTM training and were unable to form LTM **(Fig. 1A)**. These findings support a critical requirement for Bap60 in LTM.

**Figure 1.**
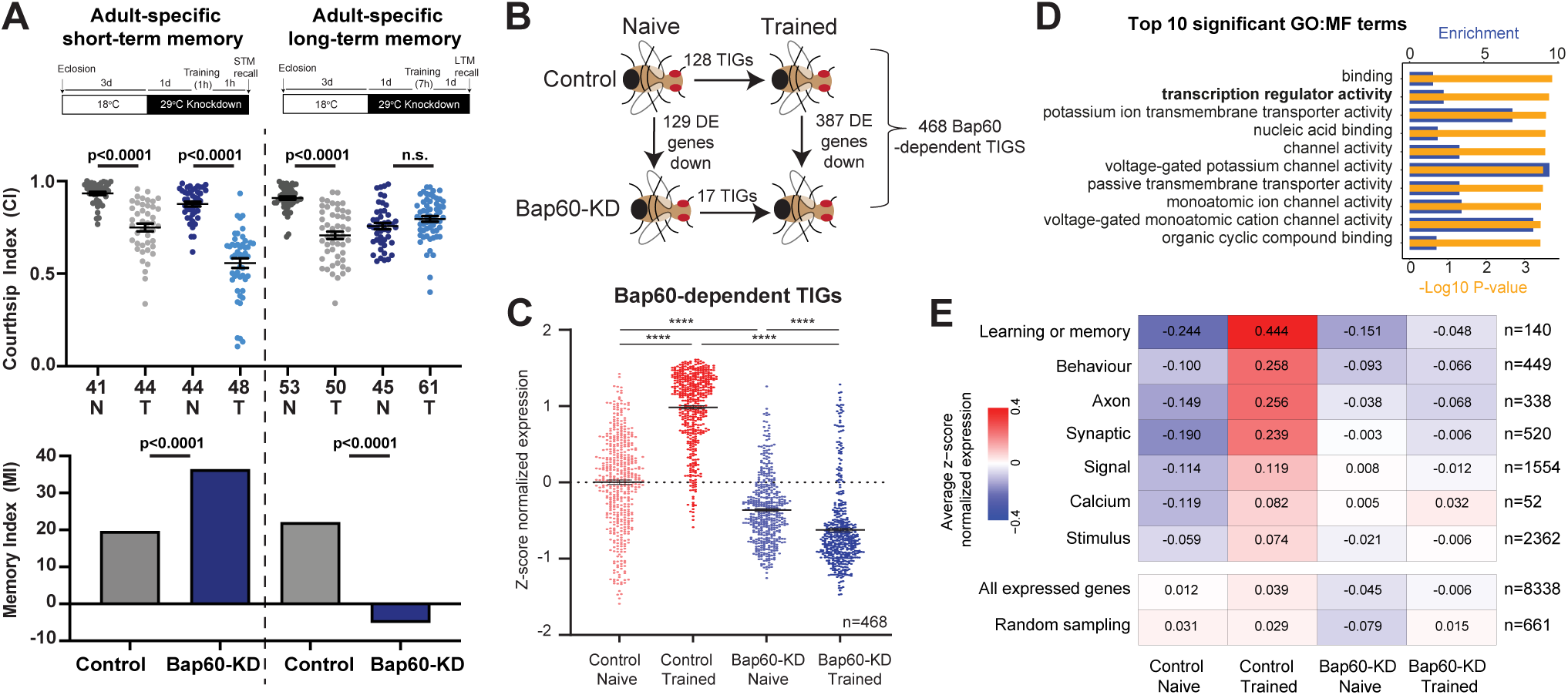
Bap60 is required for adult-specific LTM training-induced transcription. (A, top) Schematic of approach for achieving adult-specific knockdown in the MB of Bap60 (Bap60-KD). Adult-specific RNAi knockdown was facilitated by using GAL80ts, which prevents Gal4-mediated transcription at 18°C but becomes inactivated at 29°C. Knockdown of Bap60 was limited to one day prior to memory testing. Bap60-KD and genetic control flies also had expression of Unc84::GFP to enable downstream MB genomic profiling. (A, middle) Dot plot showing courtship indices (CIs) for naive (N) and trained (T) flies of Bap60-KD (blue), and its genetic control (grey), for short-term (STM, left) and long-term memory (LTM, right) assays. Statistical significance between naive and trained flies was determined using a two-tailed Mann-Whitney test. Mean is displayed with error bars indicating +/- SEM. Number of flies tested for each condition is shown under corresponding dot plot. (A, lower) Bar graph showing corresponding memory index (MI) derived from the CI (see methods). Statistical significance between MIs was determined using a randomization test with 10,000 bootstrap replicates. (B) Schematic for how Bap60-dependent training-induced genes (TIGs) were identified. Bap60-dependent training-induced genes (TIGs) were defined as those induced by training in controls but not in Bap60-KD and/or with higher post-training expression in controls compared to Bap60-KD. (C) Dot plot of z-score normalized expression values for Bap60-dependent TIGs. Genes are plotted for four different conditions: control or Bap60-KD, trained and naive. Significant differences between conditions were determined using a two-tailed pairwise t-test with Benjamini-Hochberg FDR corrected P-values indicated. (D) Bar graph of top 10 significant terms identified by gene ontology (GO) slim molecular function (MF) analysis for Bap60-dependent TIGs. Significant GO terms (FDR <0.05) were identified by contrasting Bap60-dependent TIGs to a background of all *Drosophila* genes. Enrichment (blue) and -Log10 P-values (yellow) are displayed. (E) Heatmap of average z-score normalized expression for groups of genes plotted by condition. GO terms clusters that contained memory-function associated terms including learning or memory, behavior, axon, synaptic, signal, calcium, and stimulus are shown. Z-score normalized expression scores are also shown for all expressed genes, as well as for a random set of expressed genes representing approximately 5% of coding genes in *Drosophila*.

### Bap60 is required for transcript inducibility of memory-relevant genes during LTM formation

A fundamental distinction between LTM and STM is that only LTM requires transcription. It is therefore possible that LTM training-induced transcription is compromised in Bap60-KD MBs. To investigate the role of Bap60 in transcriptional regulation during LTM formation, we isolated nuclei from the MB using INTACT)^41,46^, and performed RNA-seq on MB nuclei purified from Bap60-KD and genetic control flies one hour after LTM training and in time-of-day–matched naïve (untrained) flies (**Fig. 1B**). We then performed differential expression (DE) analysis to identify changes in expression due to training (trained vs naïve), and due to Bap60 knockdown (Control vs. Bap60-KD) (**Fig. 1B, S1A, Table S1)**. In the MB of control flies, LTM training significantly induced 128 genes when compared to naïve flies **(Fig 1B, S1A, Table S1).** In Bap60-KD flies, only 17 genes were induced after training, with eight overlapping with the control set. Also, in comparisons between Control and Bap60-KD genotypes, we found a lower number of genes misregulated under naïve conditions (129/73 genes down/up in Bap60-KD naïve) when compared to after training (387/584 genes down/up in Bap60-KD trained) (**Fig 1B, S1A, Table S1**). These results suggest that Bap60 loss has a greater impact on transcriptional responses to training than on baseline gene expression.

We defined 468 Bap60-dependent training-induced genes (TIGS) **(Fig. 1B, Table S1)** as those induced by training versus naïve in genetic controls but not in Bap60-KD, and with significantly higher post-training expression in controls than in Bap60-KD. Collectively, these TIGs were strongly induced after training in controls but showed no such induction in Bap60-KD flies, with baseline expression also being lower in Bap60-KD **(Fig. 1C).** Gene ontology (GO) enrichment analysis for molecular function revealed significant enrichment for several categories, including transcription regulator activity (GO:0140110), which was associated with 39 TIGs **(Fig. 1D, Table S2).** Because these transcriptional regulators could coordinate broader gene expression programs during LTM formation, we hypothesized that Bap60 might have a broader-ranging impact on training-induced transcription through the regulation of these transcription factors and their downstream targets; effects that may be more subtle and not fully captured through direct differential expression analysis. To test this, we analyzed transcript levels of all MB-expressed genes annotated to GO-term clusters containing key words associated with memory function, including *learning or memory*, *behavior*, *axon*, *synaptic*, *signal*, *calcium*, and *stimulus* (**Table S3**). Control flies showed clear induction across all categories, with the strongest effects observed for *learning or memory* and *behavior*. In contrast, Bap60-KD flies exhibited little to no training-induced transcription in these categories **(Fig. 1E).** Randomly sampled genes did not display training-induced expression changes in either genotype. These results demonstrate that Bap60 is either directly or indirectly required for the activity-dependent induction of a wide network of memory-relevant genes in the MB during LTM formation.

### Brm binding identifies candidate direct Bap60-dependent SWI/SNF regulated genes

As Bap60 is a core subunit of the SWI/SNF chromatin remodeling complex, we next asked whether Bap60-dependent training-induced transcriptional changes occur at genes directly bound by SWI/SNF. To address this, we profiled genome-wide binding of the SWI/SNF ATPase Brahma (Brm) in MB nuclei using Cleavage Under Targets and Release Using Nuclease (CUT&RUN). This method, like ChIP, detects protein-DNA interactions in their native chromatin context, but achieves a higher signal-to-noise ratio and works efficiently with low cell numbers, making it well suited for INTACT purified MB nuclei^47^.

To enable CUT&RUN for Brm, we obtained a FlyFos genomic construct^48^ containing a C-terminally tagged Brm protein (*2×TY1-SGFP-V5-preTEV-BLRP-3×FLAG*) from the TransgenOme project^49^. This construct was inserted the into the AttP40 landing site on chromosome 2. Expression and nuclear localization of the tagged Brm protein in the MB (and all observed brain nuclei) was confirmed by confocal microscopy **(Fig. 2A)**. MB nuclei expressing the tagged Brm protein were isolated by INTACT and subjected to CUT&RUN with antibodies against GFP and TY to detect Brm, H3K4me3 to mark active promoters, and IgG as a negative control. Brm-bound regions were defined as sites enriched over IgG in both GFP and TY datasets. This identified 6,504 Brm-bound regions corresponding to 4,388 genes **(Table S4)**. Brm binding patterns showed strong concordance with H3K4me3-marked active promoters **(Fig. 2B)**, with 3,075 Brm-bound regions also having enrichment for H3K4me3. Quantitatively, 67.6% of Brm-bound regions were located within 3kb of annotated transcription start sites (TSS) **(Fig. 2C)**, consistent with SWI/SNF occupancy at accessible, transcriptionally active promoters.

**Figure 2.**
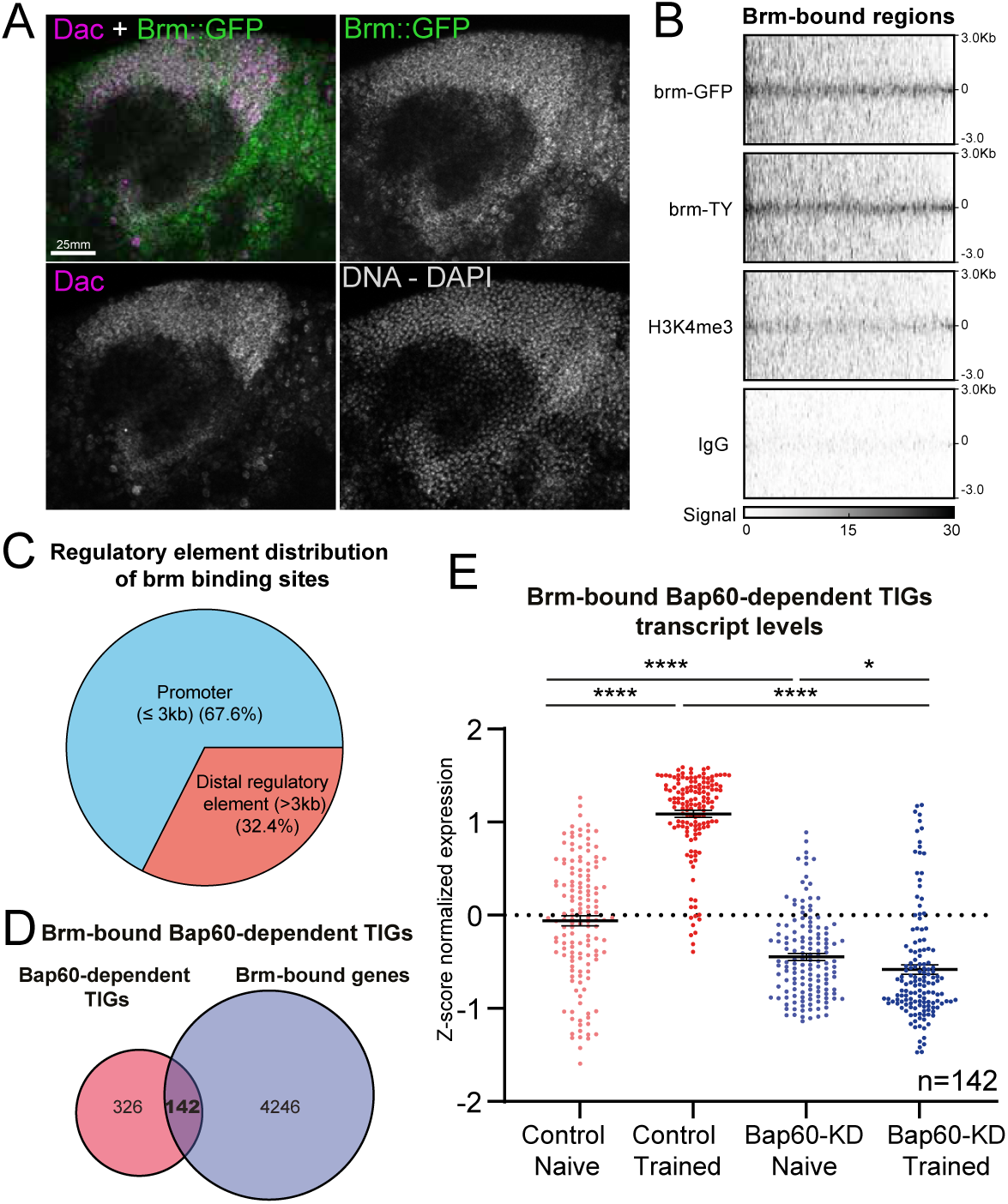
Brm CUT&RUN identifies direct SWI/SNF targets among Bap60-dependent TIGs. (A) Confocal microscopy confirming expression and nuclear localization of tagged Brm expressed from a genomic transgene in the mushroom body (MB). Brm::GFP was visualized using and anti-GFP antibody (green). Dac (magenta) was used to mark the nuclei of MB neurons. DAPI staining of DNA is shown for reference (not in overlay). (B) Heatmap of CUT&RUN signal for Brm-bound regions (n=6,504), showing concordance with H3K4me3- marked promoters and sparse IgG signal. (C) Distribution of Brm-bound regions relative to TSS: promoter (≤3 kb from TSS) versus distal regulatory element (>3 kb fro mTSS) are shown. (D) Venn diagram displaying the overlap of Brm-bound genes with Bap60-dependent training-induced genes (TIGs), with 142 shared targets (30.3%). (E) Transcript levels of Brm-bound Bap60-dependent TIGs (n=142), showing induction after training in controls but not in Bap60-KD. Significance determined by two-tailed pairwise t-test with Benjamini– Hochberg FDR correction.

We next intersected Brm-bound genes with the 468 Bap60-dependent TIGs identified in our RNA-seq analysis **(Fig. 2D)**. Of these TIGs, 142 (∼30%) were Brm-bound, representing candidate genes directly regulated by SWI/SNF during LTM formation. Transcript level analysis of the 142 Brm-bound Bap60-dependent TIGs revealed robust induction after training in control flies, but no induction in Bap60-KD flies **(Fig. 2E)**. These results indicate that Brm occupancy is associated with training-inducible expression of this subset of genes, and that Bap60 is required for their activation during LTM formation.

Notably, Brm-bound Bap60-dependent TIGs include genes with clear links to neuronal communication and plasticity. Several are involved in the regulation of neurotransmitter secretion, including the ion channel *Ih*, the active zone proteins *cpx* and *Rim*, and the membrane trafficking regulators *Syt4* and *CASK*. Others contribute to synaptic plasticity, such as *brp* and *Sap47*. Additional members participate in potassium ion transmembrane transport, including *shab*, *Atpα, Irk3*, and *Task6*. Strikingly, 24 of the Brm-bound Bap-dependent TIGs are annotated to DNA-templated transcription, encompassing transcription factors such as *crp*, *dati*, *FoxK*, *mirr*, *chinmo*, *jim*, *pros*, and *scrt*.

Together, these findings define a set of candidate direct SWI/SNF target genes whose inducible expression during LTM formation depends on Bap60, reinforcing a central role for Bap60 and the SWI/SNF complex in transcriptional activation of memory genes.

### Bap60 acts downstream of memory IEGs *Hr38* and *sr*

IEGs are rapidly induced by neuronal activity. They are often enriched for transcription factors, which can play a pivotal role in shaping downstream memory-related transcriptional programs. Upon *Bap60* knockdown in the MB, almost all training-induced gene expression was eliminated; however, 17 genes remained significantly inducible after training **(Fig.1B, S1, Table S1)**. Two of these, *Hr38* and *stripe* (*sr*), are known *Drosophila* memory IEGs factors that are robustly induced in the MB after LTM training^31,41,42^ and have been shown to be required for LTM formation and social behaviour^31,50^. In our dataset, *Hr38* transcript levels increased ∼5.7-fold and *sr* ∼2.4-fold after training in controls, with indistinguishable induction magnitudes in Bap60-KD flies **(Fig. 3A-B)**. CUT&RUN mapping revealed that Brm binds to promoter regions with MB accessible chromatin for both *Hr38* and *sr* **(Fig. 3C)**. However, although *Hr38* and *sr* are directly bound by SWI/SNF at accessible promoters, their rapid and robust induction after training is preserved independently of Bap60. This raises the possibility Bap60 may contribute to LTM training induced gene activation downstream of IEGs.

**Figure 3.**
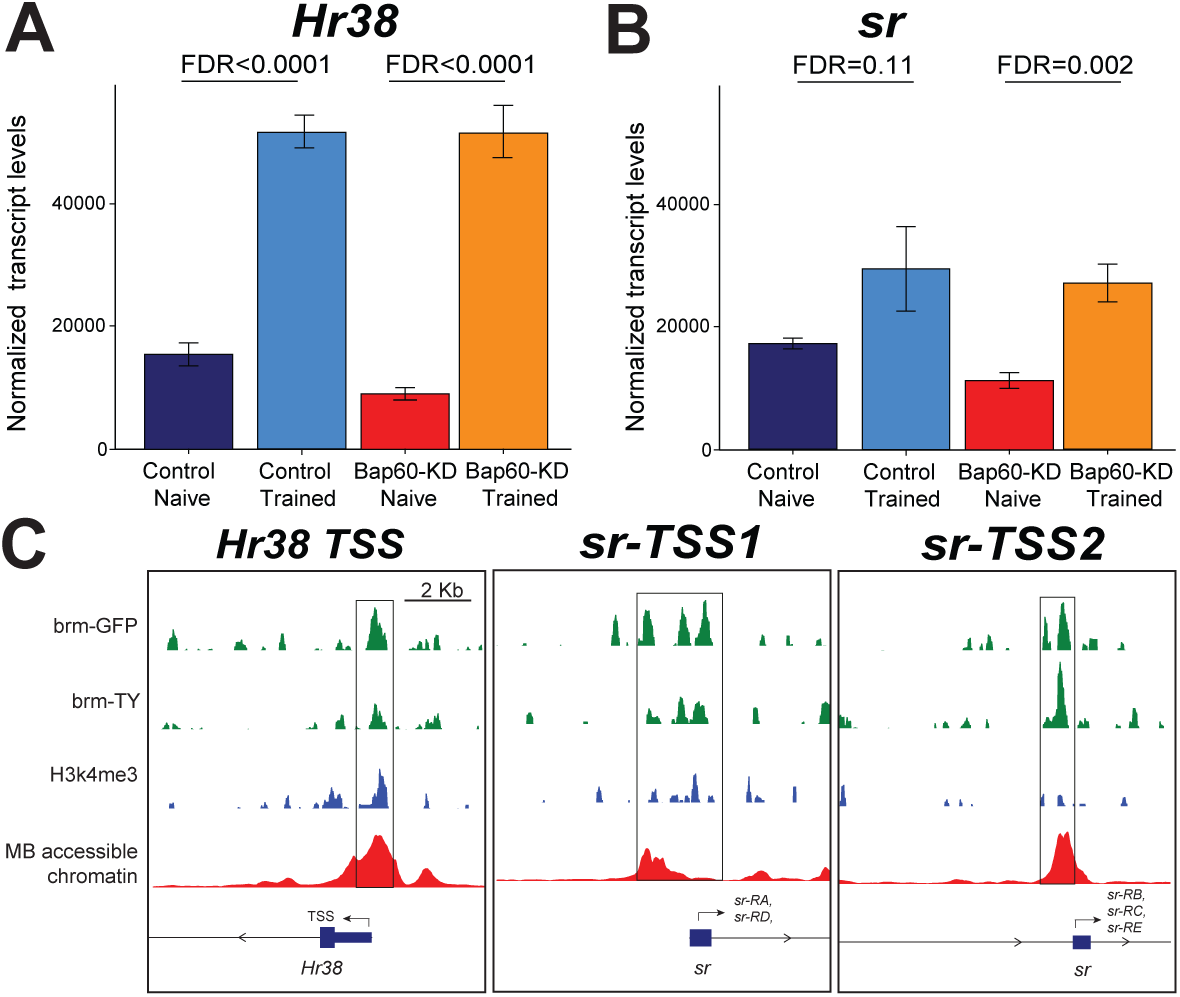
Memory IEGs Hr38 and sr are bound by Brm but do not require Bap60 for training-induced transcription. (A-B) Bar graphs of normalized MB transcript levels of courtship memory IEGs (A) *Hr38* and (B) *sr* for Bap60-KD and controls after LTM training compared to naive flies. Error bars display standard error. Differential expression analysis FDR values are shown between trained and naive conditions. (C–D) Track figures showing CUT&RUN signal at transcriptional start sites (TSS). (C) *Hr38* at the main TSS; (D) *sr* at the (left) main TSS and (right) alternate TSS. Tracks display Brm binding (Brm-GFP, Brm-TY), H3K4me3 enrichment and MB-accessible chromatin (MB ATAC-seq).CUT&RUN signals for Brm and H3K4me3 were adjusted by IgG subtraction. Grey boxes indicate regions of Brm binding, H3K4me3 signal, and MB-accessible chromatin.

To explore this hypothesis, we leveraged publicly available ChIP-seq datasets for Sr and Hr38 and overlapped these binding sites with previously generated, genetic background–matched, MB-specific ATAC-seq peaks^42^ to identify putative binding sites within accessible chromatin in adult MB neurons. Interestingly, 43 out of 142 (30%) Bap60-dependent Brm bound genes had Sr binding sites, an overlap that is significantly more than predicted by chance (p = 6.09×10^-7^, hypergeometric test) (**Fig. 4A, Table S5**). Among these 43 genes, 7 encode for DNA binding TFs, including the known MB developmental regulator Prospero (Pros)^51^. The *pros* gene contains a prominent Sr binding site that overlaps directly with Brm occupancy at a region of MB accessible chromatin enriched for H3K4me3 marks **(Fig. 4B)**. This site is located at a downstream alternative *pros* promoter controlling three isoforms (*pros-RK*, *pros-RJ*, and *pros-RL*) **(Fig. 4B)**.

**Figure 4.**
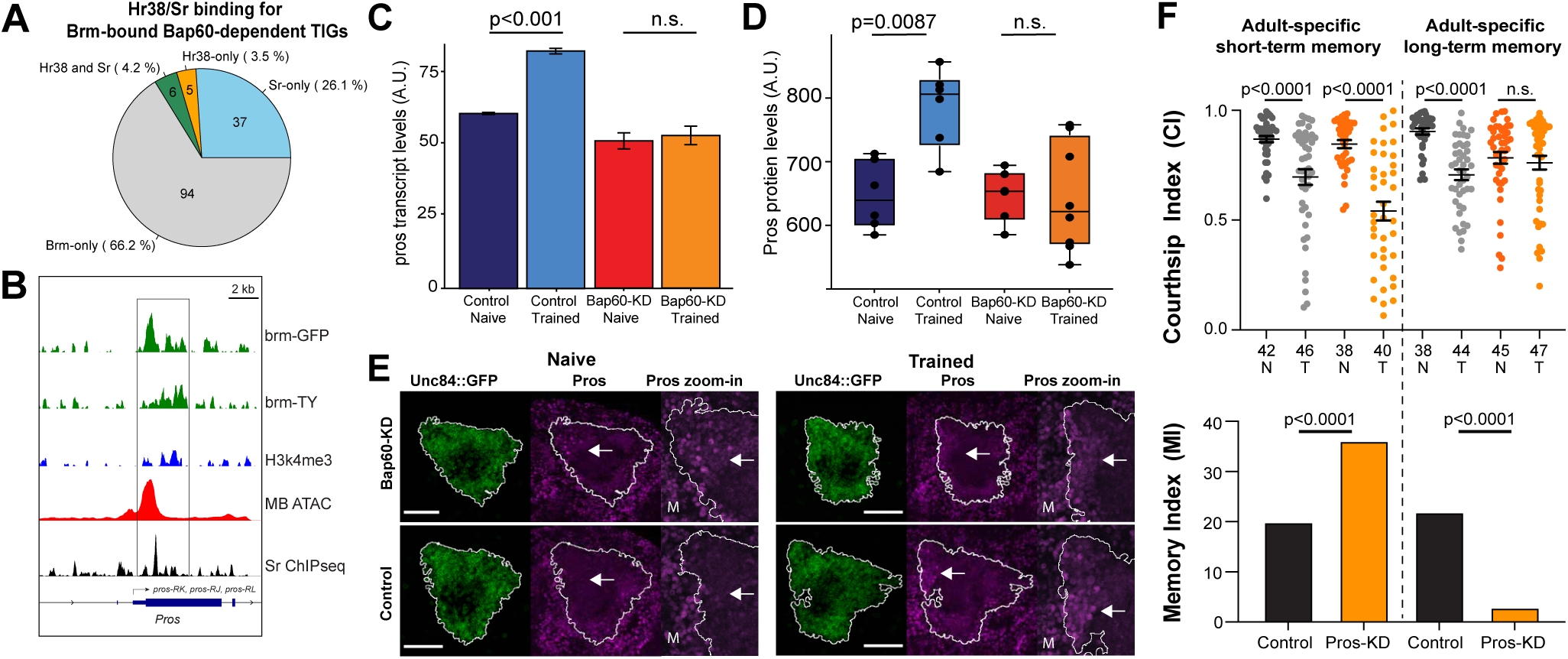
Brm-bound Bap60-dependent *Prospero* is required in the MB specifically for LTM formation. (A) Pie chart displaying the percentage of Brm-bound Bap60-dependent training-induced genes (TIGs) are bound in MB-accessible regions by Hr38 and/or Sr, as well as those only bound by Brm. (B) Track figure of the alternate transcriptional start site for *Prospero* showing CUT&RUN signal for Brm (Brm-GFP, Brm-TY), H3K4me3 enrichment, MB-accessible chromatin as determined by MB ATAC-seq, and Sr-bound chromatin from ChIPseq. CUT&RUN signal for Brm binding and H3K4me3 tracks was adjusted by subtracting IgG. Grey square indicates region of Brm-binding, H3K4me3 enrichment, MB-accessible chromatin and significant Sr-binding. (C) Bar graph showing Prospero transcript levels form RNAseq. Significance was determined using DESeq2 (Wald’s test). (D) Prospero protein levels quantified from confocal images. Quantification was performed on maximum projections of the entire region containing MB nuclei (labeled with unc84::GFP). Statistical significance was determined using a two-tailed t-test. (E) Representative confocal Z-stack max projections showing MB nuclei (unc84::GFP, green) and Prospero (anti-Pros, magenta) for naive (left), trained (right), Bap60-KD (top) and control (bottom) brains. Outline in white highlights MB nuclei and distinguishes regions where changes in Prospero fluorescence were measured. White arrow indicates the approximate location near the midline of the brain (M) where a cluster of brighter Prospero positive nuclei that often occur in trained controls. The right panel (Pros zoom in) show higher magnification views of the Prospero signal in the region indicated by the white arrow. Scale bars indicate 50 microns. (F) Dot plot showing courtship indices (CIs) for naive (N) and trained (T) flies of Prospero RNAi knockdown (Pros-KD, blue), and its control (grey), for short-term (STM, left) and long-term memory (LTM, right) assays. Statistical significance between naive and trained flies was determined using a two-tailed Mann-Whitney test. Mean is displayed with error bars indicating +/- SEM. Number of flies tested for each condition is shown under corresponding dot plot. Bar graph showing memory index (MI) derived from corresponding CI above (see methods). Statistical significance between MIs was determined using a randomization test with 10,000 bootstrap replicates.

### Prospero is required for LTM formation

Pros is a homeobox transcription factor with a well-characterized role in MB development and neuronal differentiation^52–54^. Consistent with previous findings^41^, *pros* mRNA is clearly induced in control MBs at 1 hour after training, but not in Bap60-KD MBs (**Fig. 4C**). We used an anti-pros antibody to examine Pros protein levels in the adult MB nuclei revealed similar results as mRNA with induction in controls, and not in Bap60-KD MBs (**Fig. 4D**). Interestingly, analysis of confocal images suggested that Bap-60-dependent Pros induction occurred in only a subset of MB nuclei located towards the midline of the brain (**Fig. 4E**). Fittingly, adult-specific MB RNAi knockdown of Pros one day prior to training caused phenotypes consistent with those seen in Bap60-KD flies (**Fig. 4F**). Pros-KD flies showed significantly enhanced STM compared to controls one-hour after training, but completely failed to form LTM, as evidenced by lack of courtship suppression 24 hours after training **(Fig. 4F).** These results suggest a critical role for Pros in LTM formation downstream of Bap60.

## Discussion

Mutations in components of the SWI/SNF complex in humans are a major cause of Autism Spectrum Disorders and Intellectual Disability^55,56^. Therefore, it is important to understand the functions of the SWI/SNF complex in the brain. Here, we investigated the *Drosophila* SWI/SNF subunit Bap60 in the context of LTM dependent transcriptional plasticity. We show that Bap60 is required in the MB for memory induced transcription and appears to govern a broad network of memory genes through both direct and indirect mechanisms. Bap60 was not required for LTM training induced expression of memory IEGs, *Hr38* and *sr*. However, many Bap60 dependent LTM training induced genes were co-bound by Brm and Sr including the TF Pros, which is induced by training in a subset of MB neurons and required for LTM. Interestingly, SWI/SNF^57^, and the Bap60 mouse ortholog Baf60a^58^, have been shown to physically interact with the IEG Fos/Jun TF dimer. Consistent with this, our findings support a role for Bap60 in LTM training induced transcription downstream of

### *Drosophila* memory IEGs

In *Drosophila*, an analysis of null mutants demonstrated that Bap60 is an essential gene^59^. While it does not appear to be required for core chromatin remodeling activity *in vitro*^60^, it is required for SWI/SNF mediated transcriptional activation^59^. Using targeted knockdown in the MB, we have found that Bap60 is critical at crucial developmental transitions that likely require large changes in transcription^43,44^. At the pupal stage, the MB undergoes dramatic remodeling that is initiated by a wave of transcription mediated by the Ecdysone hormone^61^. Bap60 is required for proper initiation of remodeling^44^. After pupal morphogenesis, newly eclosed flies undergo a period of experience dependent development in the MB^62,63^. We found Bap60 was required at this stage for expression of neuronal genes in the MB but had little effect in steady state flies that were 5 days old^43^. Here we observed a similar trend, with naïve Bap60-KD MBs showing modest numbers of DE genes compared to controls. However, when exposed to LTM training, Bap60 had a much larger role in LTM training induced gene expression. Overall, this highlights the importance of Bap60 for transcriptional plasticity in general, whether during development or due to environmental effects.

The mechanisms of how Bap60 mediates different types of transcriptional plasticity in flies is still unclear. Bap60 is the *Drosophila* ortholog of the mammalian SMARCD1/2/3 proteins, which are also known as Baf60a/b/c. This family of proteins contain three conserved coiled coil domains and a characteristic SWIB domain. The SMARCD family are core components of SWI/SNF that are required for its full assembly^64^. However, reconstituted minimal SWI/SNF complexes lacking SMARCD subunits still show chromatin remodeling activity in vitro^60^. SMARCD proteins with missense mutations or domain deletions still integrate into SWI/SNF complexes but are lacking critical TF interactions that mediate specific biological functions^65–67^. It will be interesting to investigate if Bap60 mediates any specific interactions between SWI/SNF and TFs that govern the memory transcriptome in *Drosophila*. Based on ChIPseq DNA binding data we hypothesize that SWI/SNF coregulates genes with Sr however, more experiments are required to determine if this is a specific interaction or an association.

## Methods

### Fly stocks and genetics

*Drosophila melanogaster* stocks were maintained on standard cornmeal-sucrose-agar medium supplemented with the mold inhibitors methyl paraben and propanoic acid, at 25 °C with 70% humidity under a 12h:12h light/dark cycle. Wild-type female flies were an in-house generated Canton-S/Oregon-R mixed line (Nijmegen wild type). *5xUAS-Unc84::2xGFP* flies were kindly provided by G.L. Henry^46^. All additional stocks were obtained from the Bloomington *Drosophila* Stock Center (BDSC; Bloomington, USA). UAS-RNAi stocks were generated by the Transgenic RNAi project (TRiP, Harvard University)^68^. *R14H06-Gal4*, which express Gal4 under the control of a MB-specific enhancer of *Adcy1* (BDSC #48677), was used to drive MB-specific expression of UAS-inducible RNAi constructs^44,69^. To facilitate CUT&RUN experiments for Brm binding, we obtained a tagged *brm* genomic clone from the TransgenOme project (Clone ID 1686682740821248 G09)^49^. This construct was created from a genomic clone from the FlyFos project -FlyFos026944^48^. This clone was inserted into the AttP40 landing site on chromosome 2 at Genome Prolab using standard methods. The resulting flies express Brm with a C-terminal *2×TY1-sGFP-V5-preTEV-BLRP-3×FLAG* tag (**Fig. 3A**).

Temporal control of Gal4-mediated RNAi knockdown was achieved using Gal80^ts^, which is expressed ubiquitously under control of the *αTub84B* promoter^70^. For Gal80^ts^ experiments, flies were raised at 18°C to prevent RNAi expression during development and shifted to 29°C one day prior to courtship LTM training to induce adult-specific knockdown. Experimental and genetic background control flies were generated using an isogenic heterozygous breeding strategy. Adult-specific Bap60 knockdown was achieved by crossing *Gal80^ts^;{UAS-Bap60^RNAi^}AttP2* (BDSC #32503) and *Gal80^ts^*;*UAS-mCherry^RNAi^* (BDSC #35785 - used as a genetic control for both background and potential off-target RNAi effects) to *5xUAS-Unc84::2xGFP*;*R14H06-Gal4*. Adult-specific Prospero knockdown was performed by crossing the TRiP RNAi line BDSC #42538 and the genetic background control BDSC #36304 to *Gal80^ts^;R14H06-Gal4*. All experimental crosses and genotypes are shown in **Table S6**.

### Courtship conditioning assay

Courtship memory was assessed using a modified version of the previously established assay^34^. Newly eclosed F1 male flies were collected and housed individually in 96-well blocks containing 500 µL of food for five days. Males were divided into naïve and trained cohorts. STM was evaluated by pairing males with unreceptive, recently mated females for two hours, followed by a one-hour isolation period. LTM was measured after a single seven-hour training session, followed by 20–24 h of isolation. After the respective rest periods, each male was paired with a new mated female, and courtship activity was recorded over 10 minutes. The courtship index (CI) was calculated as the percentage of time a male engaged in courtship behaviors over a 10-minute period. Memory performance was quantified as the memory index (MI), defined as MI = (CI_naïve_ − CI_trained_)/CI_naïve_^71^. Statistical comparisons of CIs between naïve and trained flies were performed using a two-tailed Mann-Whitney test, with outliers identified and removed via the ROUT method (GraphPad Prism v10.4.1, FDR = 1%). Differences in MI between genotypes were assessed using a two-tailed randomization test with 10,000 bootstrap replicates^34^.

### Bap60-KD sample collections

For RNA-seq experiments, F1 males carrying either adult-specific Bap60 knockdown (*Gal80^ts^*;*UAS-Bap60^RNAi^* × *5xUAS-Unc84::2xGFP*;*R14H06-Gal4*) or the corresponding mCherry^RNAi^ control (*Gal80^ts^*;*UAS-mCherry^RNAi^*× *5xUAS-Unc84::2xGFP;R14H06-Gal4*) were raised at 18 °C to suppress RNAi during development. Male flies were collected at eclosion and housed individually at 18 °C for 4 days. One day prior to LTM training, flies were shifted to 29 °C to induce knockdown and *Unc84::GFP* expression, which was confirmed by standard fluorescence microscopy. Trained males underwent a single seven-hour courtship session with unreceptive, mated females, while naïve males were collected in parallel. Approximately 50 flies per cohort were flash-frozen in liquid nitrogen one hour after training, and heads were separated from bodies using vortexing and frozen sieves. Samples were stored at −80 °C until RNA extraction and library preparation.

### Isolation of nuclei tagged in a specific cell-type (INTACT)

To obtain MB nuclei for downstream RNA-seq and CUT&RUN analyses, INTACT was performed as previously described^41,46^. Samples containing approximately 50 fly heads were homogenized in buffer containing 0.3% NP-40 using a Dounce homogenizer. The homogenate was filtered through a 40 µm mesh to remove debris. MB nuclei were then enriched by immunoprecipitation with anti-GFP antibody (Invitrogen: G10362) conjugated to magnetic beads (Invitrogen: 10004D), following the manufacturer’s instructions. The isolated nuclei were subsequently processed for RNA-seq and CUT&RUN experiments.

### RNA-sequencing and data analysis

Total RNA was extracted from immunoprecipitated MB nuclei using the Arcturus PicoPure RNA Isolation Kit (ThermoFisher Scientific, KIT0204), followed by on-column DNase digestion (Qiagen, 79254) according to the manufacturers’ protocols. RNA integrity was evaluated using the Bioanalyzer 2100 with the Pico RNA kit (Agilent, 5067-1513). cDNA synthesis and library preparation were performed with the Nugen Ovation *Drosophila* RNA-seq System 1-16 Kit. Final library size distributions and quality metrics were determined using the Bioanalyzer 2100 DNA High Sensitivity kit (Agilent, 5067-4626). Libraries were sequenced on an Illumina NovaSeq 6000 platform at Genome Québec using an S4 v1.5 200-cycle kit, generating 100 bp paired-end reads.

RNA libraries (n = 8; duplicates for each Bap60-KD and genetic controls, before and after LTM training) yielded an average of 239,898,050 reads per sample. All computational steps were run on Compute Canada servers (StdEnv/2020). First, raw reads were lightly trimmed and adapters removed with Trimmomatic (v0.39)^72^. Read quality was then assessed with FastQC (v0.11.9) and trimmed reads aligned to the *Drosophila melanogaster* reference genome (Ensembl release 103, dm6) using STAR (v2.7.5a), and retaining only uniquely aligned reads with ≤4 mismatches^73,74^. An average of 145,540,296 reads per sample mapped to annotated genes using HTSeq-count (v0.7.1) under the union mode. Low-abundance features were removed by filtering out rRNA and other non-coding RNAs, genes on the Y chromosome or mitochondrial genome, and the lowest one-third of genes by normalized expression level. This filtering step retained 8,626 robustly expressed genes for downstream analysis.

Differential expression was performed with DESeq2 (v1.30.1) in RStudio (v4.0.3)^75^, with significance determined using a two-sided Wald test, with p-values adjusted using the Benjamini-Hochberg false discovery rate (FDR). Differentially regulated genes were defined as FDR < 0.05. Gene ontology (GO) enrichment analysis was carried out using the PANTHER Molecular Function GO-Slim dataset^76^. Statistical significance of term enrichment was assessed using the one-tailed Fisher’s exact test for overrepresentation and corrected using the Benjamini-Hochberg false discovery rate (FDR). Terms were considered enriched at FDR < 0.05. Data visualization was performed using ggplot2 (v3.4.2), pheatmap (v1.0.12) and GraphPad Prism (v10.4.1). Venn diagrams were created with BioVenn^77^. Additional processing, annotation, and statistical comparisons were implemented with functions from the BinfTools package (https://github.com/kevincjnixon/BinfTools), including: count_plot, getSym, barGene, zheat, and customGMT to make custom gene sets. For the heatmap shown in Figure 2D, we generated custom gene sets based on Gene Ontology (GO) biological process terms, including ‘learning or memory’ (GO:0007611) and ‘behavior’ (GO:0007610), as well as groups of related terms for axon (e.g., GO:0007409, GO:0061564, GO:0007411, GO:0007414), synaptic processes (e.g., GO:0007268, GO:0098916, GO:0099537, GO:0099536, GO:0007416, GO:0050808, GO:0051124, GO:0050804, GO:0099177, GO:0099643, GO:0016079, GO:0045886, GO:0051964, GO:1905809, GO:0008582, GO:0099565, GO:0060079, GO:0099504, GO:0099003, GO:0001507, GO:0051963, GO:0050803, GO:0050805), signaling (e.g., GO:0007267, GO:0099177, GO:0099643, GO:0023061, GO:0023052), calcium-related processes (e.g., GO:0070509, GO:0016338, GO:0090676, GO:0070588), and stimulus responses (e.g., GO:0009605, GO:0009628, GO:0009612, GO:0050896, GO:0051602, GO:0071868, GO:0071870, GO:0009266). The full list of terms and genes used is provided in **Table S3**.

### CUT&RUN library preparation and data analysis

To determine regions of Brm binding as well as H3K4me3 enrichment, we performed Cleavage Under Targets and Release Using Nuclease (CUT&RUN) on INTACT-isolated bead-bound mushroom body (MB) nuclei. Nuclei were extracted from groups of ∼50 adult male *Drosophila* heads per replicate, each expressing *5×UAS-Unc84::2×GFP* under *R14H06-Gal4* control and 2*×TY1-SGFP-V5-preTEV-BLRP-3×FLAG*-tagged Brm. Each sample corresponded to an independent collection of ∼50 fly heads.

CUT&RUN was carried out using the CUTANA™ CUT&RUN Kit, Version 3 (EpiCypher, 14-1048) with minor modifications to accommodate INTACT-isolated nuclei, as described previously^78^. Briefly, purified bead-bound nuclei were resuspended in 100 µL of cold Antibody Buffer, and the manufacturer’s protocol was initiated at Section IV, “Antibody Binding.” For quality control, the SNAP-CUTANA K-MetStat Panel (EpiCypher) was spiked into H3K4me3 and IgG samples to assess antibody performance on designer nucleosomes while simultaneously generating data from fly nuclei. To assess Brm occupancy, 0.5 µg of antibodies against GFP (Sigma, 11814460001) and TY1 (ThermoFisher, A01004-40) were used with samples incubated overnight at 4 °C under gentle end-over-end rotation.

CUT&RUN libraries were prepared from isolated DNA using the CUTANA™ Library Prep Kit (EpiCypher,14-1001) according to the manufacturer’s instructions. Libraries were sequenced on an Illumina NovaSeq 6000 platform at Genome Québec using an S4 v1.5 200-cycle kit, generating 100 bp paired-end reads. On average, 20,529,279 reads per sample were obtained.

Raw reads were processed with Trimmomatic (v0.39) to remove low-quality reads and clip adaptor sequences. Trimmed reads were aligned to the *Drosophila melanogaster* genome (Ensembl release 103, dm6) using Bowtie2 (v2.4.1) with the parameters -x 2000 -- very-sensitive^79^. Resulting BAM files were sorted and indexed with SAMtools (v1.11)^80^. For visualization, deepTools bamCoverage was used to generate bigWig files for genome browser inspection and bedGraph files for peak calling. Regions of Brm binding and H3K4me3 enrichment over IgG signal were identified using Sparse Enrichment Analysis for CUT&RUN (SEACR) with normalization enabled (norm) and stringent mode^81^. Brm-bound regions were defined as those showing enrichment in both GFP and TY antibody samples, yielding 6,504 regions, which were annotated to 4,388 genes using the ChIPseeker R package (v1.26.2)^82^. Heatmap visualization of CUT&RUN signal over Brm-bound regions was generated with the computeMatrix and plotHeatmap functions from deepTools^83^. Genomic track visualizations for both CUT&RUN and MB-ATAC-seq datasets (GSE273839) were produced from bigWig files using pyGenomeTracks^84^. IgG signal subtracted bigwig files for track visualization of CUT&RUN datasets were produced using the bigwigCompare function from deepTools.

### ChIP-seq data analysis

To identify potential binding sites for Hr38 and Sr, publicly available ChIP-seq data were obtained from the ENCODE repository (Hr38: ENCFF144OZH; Sr: ENCFF186BCY, ENCFF247KLE)^85^. ChIP-seq peaks were generated using optimal IDR thresholding by ENCODE^86^. To enrich for sites relevant in the adult mushroom body, ChIP-seq peaks were overlapped with previously published MB-specific ATAC-seq peaks from mCherry^RNAi^ control flies (available at GEO:GSE273839)^42^. Peaks were annotated to the nearest gene using ChIPseeker (v1.26.2). Genome browser tracks were generated with pyGenomeTracks using signal p-value bigwig files generated by ENCODE.

### Staining and microscopy

Dissected brains were fixed in ice-cold 4% paraformaldehyde for 45 min at room temperature, blocked in 5% normal goat serum (NGS) for 1 h, and incubated overnight at 4 °C with primary antibodies (anti-GFP, 1:100, Invitrogen G10362; anti-Prospero, 1:20, DSHB MR1A; anti-Dac, 1/50; DSHB mAbdac1-1). Brains were then incubated overnight at 4 °C with secondary antibodies (Alexa Fluor 488, 568, and 594, 1:300, Invitrogen), mounted in SlowFade Antifade (Invitrogen: S36972), and stored at 4 °C until imaging. Confocal images were acquired on a Zeiss LSM 710 or a Leica SP8. Mean Prospero intensity was quantified in ImageJ from maximum z-stack projections within the UNC84::GFP-positive nuclei. Three to four brains were quantified for each condition and statistical significance was assessed using a two-tailed αt-test.

## Acknowledgements

We would like to thank: Genome Quebec for providing sequencing services; the Atlantic Computational Excellence Network (https://www.ace-net.ca/) and the Digital Research Alliance of Canada (alliancecan.ca) for providing access to by high performance computational infrastructure and training; the microscopy core facility at Dalhousie University for access to confocal microscopes; the TransgenOme and FlyFos projects based at the Max Plank Institute of Molecular Cell Biology and Genetics for creating and distributing the tagged brm genomic construct; the Bloomington *Drosophila* Stock Center at Indiana University, the Vienna *Drosophila* Resource Center (VDRC), and Dr. G.L. Henry for fly stocks used in this study, and the MODERN resource and ENCODE for providing publicly available ChIP-seq data. Finally, we would like thank Dr. Kevin Nixon for providing his expert guidance and coding knowledge to help with our bioinformatic analysis, and Robert Reid-Taylor for his continuous aid to operations in the Kramer lab.

## Data availability

Raw and processed RNA-seq and CUT&RUN data are submitted to the GEO database under the accession number GSE305647.

## Author contributions

JMK and SGJ conceptualized and designed the project; SGJ performed the courtship memory experiments; SGJ performed the INTACT RNA-seq and CUT&RUN experiments and analyzed the resulting data; NR performed the Prospero confocal microscopy and analysis; SB and AH performed confocal imaging of the GFP-tagged Brm transgenic fly line; SGJ and JMK wrote the original draft; SGJ, NR and JMK made the figures; JMK supervised the project and was responsible for funding acquisition.

## References

1. Alberini, C. M. & Kandel, E. R. The Regulation of Transcription in Memory Consolidation. Cold Spring Harb. Perspect. Biol. 7, a021741 (2015).

2. Chen, X., Rahman, R., Guo, F. & Rosbash, M. Genome-wide identification of neuronal activity-regulated genes in drosophila. Elife 5, 1–21 (2016).

3. Yap, E. & Greenberg, M. E. Review Activity-Regulated Transcription : Bridging the Gap between Neural Activity and Behavior. Neuron 100, 330–348 (2018).

4. Pittenger, C. & Kandel, E. R. In search of general mechanisms for long-lasting plasticity: Aplysia and the hippocampus. Philos. Trans. R. Soc. Lond. B. Biol. Sci. 358, 757–63 (2003).

5. Yin, J. C. P. et al. Induction of a dominant negative CREB transgene specifically blocks long-term memory in Drosophila. Cell 79, 49–58 (1994).

6. Dudai, Y., Jan, Y. N., Byers, D., Quinn, W. G. & Benzer, S. dunce, a mutant of Drosophila deficient in learning. Proc. Natl. Acad. Sci. U. S. A. 73, 1684–8 (1976).

7. Montarolo, P. G. et al. A critical period for macromolecular synthesis in long-term heterosynaptic facilitation in Aplysia. Science 234, 1249–54 (1986).

8. Brunelli, M., Castellucci, V. & Kandel, E. R. Synaptic facilitation and behavioral sensitization in Aplysia: possible role of serotonin and cyclic AMP. Science 194, 1178– 81 (1976).

9. Kaldun, J. C. & Sprecher, S. G. Initiated by CREB: Resolving Gene Regulatory Programs in Learning and Memory. BioEssays 41, 1900045 (2019).

10. Frank, D. A. & Greenberg, M. E. CREB: A mediator of long-term memory from mollusks to mammals. Cell 79, 5–8 (1994).

11. Tully, T., Preat, T., Boynton, S. C. & Del Vecchio, M. Genetic dissection of consolidated memory in Drosophila. Cell 79, 35–47 (1994).

12. Davis, R. L. Learning and memory using Drosophila melanogaster : a focus on advances made in the fifth decade of research. Genetics 224, (2023).

13. Neigeborn, L. & Carlson, M. Genes affecting the regulation of SUC2 gene expression by glucose repression in Saccharomyces cerevisiae. Genetics 108, 845–58 (1984).

14. Stern, M., Jensen, R. & Herskowitz, I. Five SWI genes are required for expression of the HO gene in yeast. J. Mol. Biol. 178, 853–868 (1984).

15. Son, E. Y. & Crabtree, G. R. The role of BAF (mSWI/SNF) complexes in mammalian neural development. Am. J. Med. Genet. Part C Semin. Med. Genet. 166, 333–349 (2014).

16. Jakub, T., Quesnel, K., Keung, C., Bérubé, N. G. & Kramer, J. M. Epigenetics in intellectual disability. in Epigenetics in Psychiatry 489–517 (Elsevier, 2021). doi:10.1016/B978-0-12-823577-5.00030-1.

17. Clapier, C. R. & Cairns, B. R. The Biology of Chromatin Remodeling Complexes. Annu. Rev. Biochem. 78, 273–304 (2009).

18. Yudkovsky, N., Logie, C., Hahn, S. & Peterson, C. L. Recruitment of the SWI/SNF chromatin remodeling complex by transcriptional activators. Genes Dev. 13, 2369– 2374 (1999).

19. López, A. J. & Wood, M. A. Role of nucleosome remodeling in neurodevelopmental and intellectual disability disorders. Front. Behav. Neurosci. 9, (2015).

20. Vogel-Ciernia, A. et al. The neuron-specific chromatin regulatory subunit BAF53b is necessary for synaptic plasticity and memory. Nat. Neurosci. 16, 552–561 (2013).

21. Lessard, J. et al. An Essential Switch in Subunit Composition of a Chromatin Remodeling Complex during Neural Development. Neuron 55, 201–215 (2007).

22. Narayanan, R. & Tuoc, T. C. Roles of chromatin remodeling BAF complex in neural differentiation and reprogramming. Cell Tissue Res. 356, 575–584 (2014).

23. Narayanan, R. et al. Loss of BAF (mSWI/SNF) Complexes Causes Global Transcriptional and Chromatin State Changes in Forebrain Development. Cell Rep. 13, 1842–1854 (2015).

24. Olave, I., Wang, W., Xue, Y., Kuo, A. & Crabtree, G. R. Identification of a polymorphic, neuron-specific chromatin remodeling complex. Genes Dev. 16, 2509–2517 (2002).

25. Tuoc, T. et al. Ablation of BAF170 in Developing and Postnatal Dentate Gyrus Affects Neural Stem Cell Proliferation, Differentiation, and Learning. Mol. Neurobiol. 54, 4618–4635 (2017).

26. Hargreaves, D. C. & Crabtree, G. R. ATP-dependent chromatin remodeling: genetics, genomics and mechanisms. Cell Res. 21, 396–420 (2011).

27. Wu, J. I. et al. Regulation of Dendritic Development by Neuron-Specific Chromatin Remodeling Complexes. Neuron 56, 94–108 (2007).

28. Rowland, M. E., Jajarmi, J. M., Osborne, T. S. M. & Ciernia, A. V. Insights Into the Emerging Role of Baf53b in Autism Spectrum Disorder. Front. Mol. Neurosci. 15, 805158 (2022).

29. Lessard, J. et al. An essential switch in subunit composition of a chromatin remodeling complex during neural development. Neuron 55, 201–15 (2007).

30. Staahl, B. T. et al. Kinetic analysis of npBAF to nBAF switching reveals exchange of SS18 with CREST and integration with neural developmental pathways. J. Neurosci. 33, 10348–61 (2013).

31. Martin, F. et al. A memory transcriptome time course reveals essential long-term memory transcription factors. (2025) doi:10.21203/rs.3.rs-5746840/v1.

32. Siegel, R. W. & Hall, J. C. Conditioned responses in courtship behavior of normal and mutant Drosophila. Proc. Natl. Acad. Sci. U. S. A. 76, 3430–3434 (1979).

33. Raun, N., Jones, S. & Kramer, J. M. Conditioned courtship suppression in Drosophila melanogaster. J. Neurogenet. 0, 1–27 (2021).

34. Koemans, T. S. et al. Drosophila Courtship Conditioning As a Measure of Learning and Memory. J. Vis. Exp. 1–11 (2017) doi:10.3791/55808.

35. Gil-Martí, B. et al. A simplified courtship conditioning protocol to test learning and memory in Drosophila. STAR Protoc. 4, (2023).

36. McBride, S. M.. et al. Mushroom Body Ablation Impairs Short-Term Memory and Long-Term Memory of Courtship Conditioning in Drosophila melanogaster. Neuron 24, 967–977 (1999).

37. Han, P.-L., Levin, L. R., Reed, R. R. & Davis, R. L. Preferential expression of the drosophila rutabaga gene in mushroom bodies, neural centers for learning in insects. Neuron 9, 619–627 (1992).

38. Aso, Y. et al. The neuronal architecture of the mushroom body provides a logic for associative learning. Elife 3, e04577 (2014).

39. Heisenberg, M., Borst, A., Wagner, S. & Byers, D. Drosophila Mushroom Body Mutants are Deficient in Olfactory Learning. J. Neurogenet. 2, 1–30 (1985).

40. de Belle, J. S. & Heisenberg, M. Associative odor learning in Drosophila abolished by chemical ablation of mushroom bodies. Science vol. 263 692–695 (1994).

41. Jones, S. G., Nixon, K. C. J., Chubak, M. C. & Kramer, J. M. Mushroom Body Specific Transcriptome Analysis Reveals Dynamic Regulation of Learning and Memory Genes After Acquisition of Long-Term Courtship Memory in Drosophila. Genes/Genomes/Genetics 8, 3433–3446 (2018).

42. Raun, N. et al. Trithorax regulates long-term memory in Drosophila through epigenetic maintenance of mushroom body metabolic state and translation capacity. PLOS Biol. 23, e3003004 (2025).

43. Nixon, K. C. J. et al. A Syndromic Neurodevelopmental Disorder Caused by Mutations in SMARCD1, a Core SWI / SNF Subunit Needed for Context-Dependent Neuronal Gene Regulation in Flies. Am. J. Hum. Genet. 104, 596–610 (2019).

44. Chubak, M. C. et al. Individual components of the SWI/SNF chromatin remodelling complex have distinct roles in memory neurons of the Drosophila mushroom body. Dis. Model. Mech. 12, (2019).

45. Chubak, M. C. et al. Systematic functional characterization of the intellectual disability-associated SWI/SNF complex reveals distinct roles for the BAP and PBAP complexes in post-mitotic memory forming neurons of the Drosophila mushroom body. BioRxiv (2018).

46. Henry, G. L., Davis, F. P., Picard, S. & Eddy, S. R. Cell type-specific genomics of Drosophila neurons. Nucleic Acids Res. 40, 9691–704 (2012).

47. Skene, P. J. & Henikoff, S. An efficient targeted nuclease strategy for high-resolution mapping of DNA binding sites. Elife 6, 1–35 (2017).

48. Ejsmont, R. K., Bogdanzaliewa, M., Lipinski, K. A. & Tomancak, P. Production of Fosmid Genomic Libraries Optimized for Liquid Culture Recombineering and Cross-Species Transgenesis. in 423–443 (2012). doi:10.1007/978-1-61779-228-1_25.

49. Sarov, M. et al. A genome-wide resource for the analysis of protein localisation in Drosophila. Elife 5, 1–38 (2016).

50. Agrawal, P., Chung, P., Heberlein, U. & Kent, C. Enabling cell-type-specific behavioral epigenetics in Drosophila : a modified high-yield INTACT method reveals the impact of social environment on the epigenetic landscape in dopaminergic neurons. BMC Biol. 17, 1–19 (2019).

51. Hatch, H. A. M., Belalcazar, H. M., Marshall, O. J. & Secombe, J. A kdm5–prospero transcriptional axis functions during early neurodevelopment to regulate mushroom body formation. Elife 10, 1–29 (2021).

52. Vaessin, H. et al. prospero is expressed in neuronal precursors and encodes a nuclear protein that is involved in the control of axonal outgrowth in Drosophila. Cell 67, 941– 953 (1991).

53. Elsir, T., Smits, A., Lindström, M. S. & Nistér, M. Transcription factor PROX1: Its role in development and cancer. Cancer Metastasis Rev. 31, 793–805 (2012).

54. Hatch, H. A., Belalcazar, H. M., Marshall, O. J. & Secombe, J. A KDM5–Prospero transcriptional axis functions during early neurodevelopment to regulate mushroom body formation. Elife 10, (2021).

55. Valencia, A. M. et al. Landscape of mSWI/SNF chromatin remodeling complex perturbations in neurodevelopmental disorders. Nat. Genet. 55, 1400–1412 (2023).

56. Cenik, B. K. & Shilatifard, A. COMPASS and SWI/SNF complexes in development and disease. Nat. Rev. Genet. 22, 38–58 (2021).

57. Vierbuchen, T. et al. AP-1 Transcription Factors and the BAF Complex Mediate Signal-Dependent Enhancer Selection. Mol. Cell 68, 1134–1146.e6 (2017).

58. Ito, T. et al. Identification of SWI·SNF Complex Subunit BAF60a as a Determinant of the Transactivation Potential of Fos/Jun Dimers. J. Biol. Chem. 276, 2852–2857 (2001).

59. Möller, A., Avila, F. W., Erickson, J. W. & Jäckle, H. Drosophila BAP60 is an essential component of the Brahma complex, required for gene activation and repression. J. Mol. Biol. 352, 329–37 (2005).

60. Ahmad, K. & Spens, A. E. Separate Polycomb Response Elements control chromatin state and activation of the vestigial gene. PLoS Genet. 15, 1–21 (2019).

61. Alyagor, I. et al. Combining Developmental and Perturbation-Seq Uncovers Transcriptional Modules Orchestrating Neuronal Remodeling. Dev. Cell 47, 38–52.e6 (2018).

62. Barth, M. & Heisenberg, M. Vision affects mushroom bodies and central complex in Drosophila melanogaster. Learn. Mem. 4, 219–29 (1997).

63. Balling, A., Technau, G. M. & Heisenberg, M. Are the structural changes in adult Drosophila mushroom bodies memory traces? Studies on biochemical learning mutants. J. Neurogenet. 4, 65–73 (1987).

64. Mashtalir, N. et al. Modular Organization and Assembly of SWI/SNF Family Chromatin Remodeling Complexes. Cell 1–17 (2018) doi:10.1016/j.cell.2018.09.032.

65. Debril, M.-B. et al. Transcription factors and nuclear receptors interact with the SWI/SNF complex through the BAF60c subunit. J. Biol. Chem. 279, 16677–86 (2004).

66. Priam, P. et al. SMARCD2 subunit of SWI/SNF chromatin-remodeling complexes mediates granulopoiesis through a CEBPɛ dependent mechanism. Nat. Genet. 49, 753–764 (2017).

67. Priam, P. et al. Smarcd1 subunit of SWI/SNF chromatin-remodeling complexes collaborates with E2a to promote murine lymphoid specification. Dev. Cell 59, 3124–3140.e8 (2024).

68. Perkins, L. A. et al. The Transgenic RNAi Project at Harvard Medical School: Resources and Validation. Genetics 201, 843–852 (2015).

69. Jenett, A. et al. A GAL4-Driver Line Resource for Drosophila Neurobiology. Cell Rep. 2, 991–1001 (2012).

70. Mcguire, S. E., Le, P. T., Osborn, A. J., Matsumoto, K. & Davis, R. L. Spatiotemporal Rescue of Memory Dysfunction in Drosophila. Science (80-.). 302, 1765–1769 (2003).

71. Keleman, K., Krüttner, S., Alenius, M. & Dickson, B. J. Function of the Drosophila CPEB protein Orb2 in long-term courtship memory. Nat. Neurosci. 10, 1587–93 (2007).

72. Bolger, A. M., Lohse, M. & Usadel, B. Genome analysis Trimmomatic : a flexible trimmer for Illumina sequence data. Bioinformatics 30, 2114–2120 (2014).

73. Aken, B. L. et al. The Ensembl Gene Annotation System. Database (Oxford). 2016, baw093 (2016).

74. Dobin, A. et al. STAR: Ultrafast universal RNA-seq aligner. Bioinformatics 29, 15–21 (2013).

75. Love, M. I., Huber, W. & Anders, S. Moderated estimation of fold change and dispersion for RNA-seq data with DESeq2. Genome Biol. 15, 1–21 (2014).

76. Aleksander, S. A. et al. The Gene Ontology knowledgebase in 2023. Genetics 224, (2023).

77. Hulsen, T., de Vlieg, J. & Alkema, W. BioVenn - a web application for the comparison and visualization of biological lists using area-proportional Venn diagrams. BMC Genomics 9, 488 (2008).

78. Jauregui-Lozano, J., Bakhle, K. & Weake, V. M. In vivo tissue-specific chromatin profiling in Drosophila melanogaster using GFP-tagged nuclei. Genetics 218, (2021).

79. Langmead, B. & Salzberg, S. L. Fast gapped-read alignment with Bowtie 2. 9, 357– 360 (2012).

80. Li, H. et al. The Sequence Alignment / Map format and SAMtools. 25, 2078–2079 (2009).

81. Meers, M. P., Tenenbaum, D. & Henikoff, S. Peak calling by Sparse Enrichment Analysis for CUT&RUN chromatin profiling. Epigenetics Chromatin 12, 42 (2019).

82. Wang, Q. et al. Exploring Epigenomic Datasets by ChIPseeker. Curr. Protoc. 2, (2022).

83. Ram, F. et al. deepTools2 : a next generation web server for deep-sequencing data analysis. 44, 160–165 (2016).

84. Lopez-Delisle, L. et al. pyGenomeTracks: reproducible plots for multivariate genomic datasets. Bioinformatics 37, 422–423 (2021).

85. Luo, Y. et al. New developments on the Encyclopedia of DNA Elements (ENCODE) data portal. Nucleic Acids Res. 48, D882–D889 (2020).

86. Landt, S. G. et al. ChIP-seq guidelines and practices of the ENCODE and modENCODE consortia. Genome Res. 22, 1813–1831 (2012).

