## Supplementary material for "The SWI/SNF complex sub-unit Bap60 is required for training-induced gene transcription during long-term memory formation": Figure S1

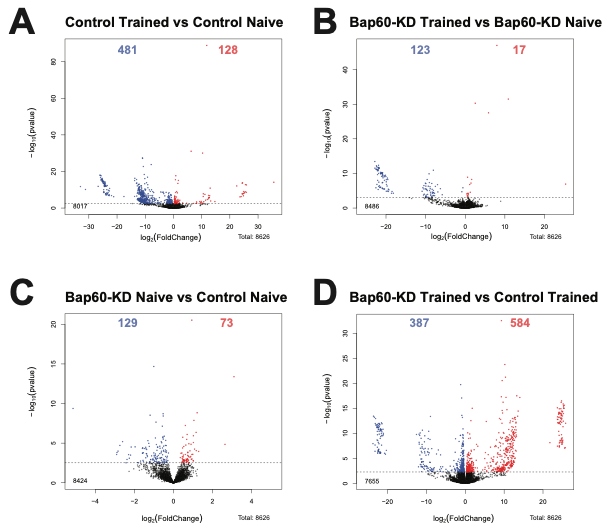
**Figure S1. Identifying Bap60-dependent training-regulated transcript level changes**

Differential expression analysis results visualized as a volcano plot for INTACT-isolated mushroom body (MB) nuclei transcript level changes between (A) control LTM vs control naive (B) Bap60-KD LTM vs Bap60-KD naive (C) Bap60-KD naive vs control naive (D) Bap60-KD LTM

vs control LTM. 8626 genes were used for differential expression analysis and declared significant if FDR < 0.05. Training induced genes are highlighted in red, and memory repressed genes are highlighted in blue.
